# Amphistomy increases leaf photosynthesis more in coastal than montane plants of Hawaiian ‘ilima (*Sida fallax*)

**DOI:** 10.1101/2023.11.03.565529

**Authors:** Genevieve Triplett, Thomas N. Buckley, Christopher D. Muir

## Abstract

**Premise of the study:** The adaptive significance of stomata on both upper and lower leaf surfaces, called amphistomy, is unresolved. A widespread association between amphistomy and open, sunny habitats suggests the adaptive benefit of amphistomy may be greatest in these contexts, but this hypothesis has not been tested experimentally. Understanding why amphistomy evolves can inform its potential as a target for crop improvement and paleoenvironment reconstruction.

**Methods:** We developed a new method to quantify “amphistomy advantage”, AA, as the log-ratio of photosynthesis in an amphistomatous leaf to that of the same leaf but with gas exchange blocked through the upper (adaxial) surface, which we term “pseudohypostomy”. We used humidity to modulate stomatal conductance and thus compare photosynthetic rates at the same total stomatal conductance. We estimated AA and related physiological and anatomical traits in 12 populations, six coastal (open, sunny) and six montane (closed, shaded), of the indigenous Hawaiian species ‘ilima (*Sida fallax*).

**Key results:** Coastal ‘ilima leaves benefit 4.04 times more from amphistomy compared to their montane counterparts. Our evidence was equivocal with respect to two hypotheses – that coastal leaves benefit more because 1) they are thicker and therefore have lower CO_2_ conductance through the internal airspace, and 2) that they benefit more because they have similar conductance on each surface, as opposed to most of the conductance being on the lower (abaxial) surface.

**Conclusions:** This is the first direct experimental evidence that amphistomy *per se* increases photosynthesis, consistent with the hypothesis that parallel pathways through upper and lower mesophyll increase the supply of CO_2_ to chloroplasts. The prevalence of amphistomatous leaves in open, sunny habitats can partially be explained the increased benefit of amphistomy in ‘sun’ leaves, but the mechanistic basis of this observation is an area for future research.

## INTRODUCTION

Amphistomy, the presence of stomata on both lower and upper surfaces of broad leaves, should increase carbon gain by reducing the average diffusion pathlength between stomata and chloroplasts, yet paradoxically this seemingly simple adaptation is uncommon in nature and we don’t know why. Understanding variation in stomatal traits like amphistomy is imperative because these tiny pores play an outsized ecological role in the global carbon and water cycles (Hetherington and Woodward, 2003; Berry et al., 2010). A widely applicable, accurate representation of how stomata mediate the relationship between CO_2_ gained through photosynthesis and water lost through transpiration is essential to predict future climate using Earth Systems Models (Jarvis, 1976; Ball et al., 1987; Collatz et al., 1991; Leuning, 1995; Sellers et al., 1997). Optimality models accurately predict the major cause of water loss, stomatal conductance (*g*_sw_), by assuming plants maximize carbon gain minus a cost of water (Cowan and Farquhar, 1977; Givnish, 1986; Medlyn et al., 2011; Lin et al., 2015; Wang et al., 2017; Franks et al., 2018; Deans et al., 2020; Franklin et al., 2020; Wang et al., 2020; Harrison et al., 2021). Despite the success of optimality modeling in predicting *g*_sw_, the same modeling approach has so far failed to explain the rarity of amphistomatous leaves (Muir, 2019). **This gap between theory and observations strongly implies that we remain ignorant about some key benefits and costs associated with stomata**.

Where are amphistomatous leaves found and why aren’t they more common? Among terrestrial flowering plants, amphistomatous leaves are rarely found on woody plants and shade-tolerant herbs, but they are common in annual and perennial herbs from sunny habitats (Salisbury, 1928; Parkhurst, 1978; Mott et al., 1982; Peat and Fitter, 1994; Gibson, 1996; Jordan et al., 2014; Muir, 2015, 2018; Bucher et al., 2017). Even in resupinate leaves where the abaxial surface faces up toward the sky, stomata develop on the lower adaxial surface (Lyshede, 2002). Exceptions to this general pattern include some arid woody plants which typically have vertically oriented, isobilateral leaves (Wood, 1934; Jordan et al., 2014; Boer et al., 2016; Drake et al., 2019) and floating/amphibious leaves of aquatic plants (Kaul, 1976; Doll et al., 2021). The dearth of amphistomatous leaves should be quite surprising and has been described as one of the most important unsolved problems in the study of leaf structure-function relations despite some recent progress (Grubb, 1977, 2020).

Amphistomatous leaves should be common because, all else being equal, a leaf with a given number of stomata per area could increase its photosynthetic rate simply by apportioning approximately half its stomata to each surface (Parkhurst, 1978; Gutschick, 1984a, b). The key difference between a hypo- and amphistomatous leaf, holding all other factors constant, is that an amphistomatous leaf has two parallel diffusion paths through the internal airspace to any given chloroplast. Those airspaces pose a resistance for CO_2_ diffusion, so CO_2_ concentration drops as it approaches chloroplasts. Shorter pathways mean a smaller drop in CO_2_ concentration. Thus, chloroplasts in amphistomatous leaves experience higher CO_2_ concentrations than in hypostomatous leaves, thereby increasing photosynthesis. The airspace resistance (or its inverse, the airspace conductance, *g*_ias_) is rarely measured directly and there is disagreement between empirical (Parkhurst and Mott, 1990; Morison et al., 2005; Evans et al., 2009; Tomás et al., 2013; Earles et al., 2018; Šantrůček et al., 2019; Nobel, 2020; Harwood et al., 2021; Márquez et al., 2023) and theoretical models (Tholen and Zhu, 2011; Ho et al., 2016; Théroux-Rancourt et al., 2021). The *g*_ias_ in thin, porous leaves may be so large as to be inconsequential given much lower conductances for other components of the diffusion pathway, whereas the *g*_ias_ of thick leaves with little airspace may greatly hinder CO_2_ diffusion to chloroplasts. Amphistomy should confer the largest photosynthetic benefit in leaves with intrinsically low *g*_ias_. The airspace conductance is one component of the overall mesophyll conductance, *g*_m_, which is often strongly influenced by the chloroplast surface area exposed to airspace and mesophyll cell wall thickness (Evans et al., 2009; Gago et al., 2020; Flexas et al., 2021). Hence, thicker leaves may compensate for lower *g*_ias_ through increased chloroplast surface area exposed to airspace (Terashima et al., 2006), but will still benefit from amphistomy as long as *g*_ias_ is finite.

Amphistomy should also enhance photosynthesis when leaf boundary layer resistance is high, because apportioning total flux between two boundary layers rather than one results in a smaller CO_2_ concentration drop between the atmosphere and stomata. A similar effect has been validated with a computer model and measurements for transpiration: amphistomatous leaves lose somewhat more water for the same vapor pressure deficit and total *g*_sw_ (Foster and Smith, 1986), but the additional carbon gain should be enough to offset this cost under most realistic conditions (Muir, 2019). However, if minimal stomatal conductance is related to stomatal density (Drake et al., 2013; Márquez et al., 2022) and the upper boundary layer conductance is higher, then amphistomy could cause additional, unavoidable water loss.

The most promising adaptive hypothesis is that amphistomy is important for maximizing photosynthetic rate under high light. Mott et al. (1982) proposed that “plants with a high photosynthetic capacity, living in full-sun environments, and experiencing rapidly fluctuating or continuously available soil water” would benefit most, in terms of increased carbon gain, from having amphistomatous leaves. As described above, herbs from sunny habitats are often amphistomatous. Most variation in stomatal density ratio (SR, the ratio of stomatal density between the upper and lower surfaces) among species is assumed to be genetic, but there is also putatively adaptive plasticity in response to light. Leaves of *Ambrosia cordifolia*, a desert perennial herb, are hypostomatous under low light (photosynthetic photon flux density, PPFD = 110 *μ*mol m^−2^ s^−1^) but develop ∼20% of their stomata on the upper surface under high light (1700 *μ*mol m^−2^ s^−1^) (Mott and Michaelson, 1991). Similarly, *Solanum lycopersicum* leaves are hypostomatous when grown in the shade but develop ∼20% of their stomata on the upper surface grown under high light-intensity (Gay and Hurd, 1975). Adult leaves of *Eucalyptus globulus* are amphistomatous, but the proportion of adaxial stomata increases from ∼10-20% under low light to ∼30-40% under high light (James and Bell, 2001). In summary, both genetic and plastic responses evince a widespread association between light and SR.

The association between high light and amphistomy suggests that ‘sun’ leaves have the most to gain in terms of increased photosynthesis from having stomata on both surfaces, as Mott et al. (1982) hypothesized. Parkhurst (1978) proposed quantifying this benefit as ‘amphistomy advantage’ (AA), which we adopt here with some modification (see Materials and Methods). This hypothesis has never been tested directly by comparing the photosynthetic rate of an amphistomatous leaf to that of an otherwise identical hypostomatous leaf with the same total stomatal conductance under the same conditions. We propose a straightforward method to do this by experimentally creating a pseudohypostomatous leaf with gas exchange blocked through the upper surface (see Materials and Methods). We use humidity to modulate stomatal conductance so that amphi- and pseudohypostomatous leaves can be compared at the same total stomatal conductance. One reason that sun leaves might have greater AA is that they are usually thicker or denser (Poorter et al., 2019), which will often result in lower *g*_ias_ either by increasing the diffusion path length (Parkhurst, 1978) or making the airspace less porous. A nonmutually exclusive hypothesis is that if sun leaves have a stomatal density ratio closer to 0.5 (same density on each leaf surface), this will confer a greater advantage than an amphistomatous leaf with most stomata on one surface. In other words, amphistomy doesn’t make much difference if one leaf surface has few open stomata on it. We therefore predict that sun leaves will have greater AA possibly because they have thicker leaves and/or SR closer to 0.5. We actually report *g*_smax,ratio_, which is similar to SR except that it accounts for differences in both stomatal density and size between surfaces.

The native flora of the Hawaiian archipelago is a excellent system to test the relationship between light habitat and AA. Many lineages have adapted to different light habitats after colonization and leaf anatomical traits such as SR and thickness vary within and among closely related species. It is hypothesized that the common ancestor in many Hawaiian clades was a weedy species with high dispersal ability adapted to open habitats (Carlquist, 1966). Colonization was followed by adaptive radiation into higher elevation, montane, closed, forested habitats. Consequently, adaptation to sun and shade is a common axis of phenotypic variation among Hawaiian plants such as lobeliads (Givnish et al., 2004; Montgomery and Givnish, 2008; Givnish et al., 2009; Givnish and Montgomery, 2014; Scoffoni et al., 2015), *Bidens* (Carlquist, 1966; Knope et al., 2020), *Scaevola* (Robichaux and Pearcy, 1984; McKown et al., 2016), *Euphorbia* (Sporck, 2011), and *Plantago* (Dunbar-Co et al., 2009).

Here we focus on variation within an indigenous plant species *Sida fallax* Walp. (Malvaceae), known in the Hawaiian language as ‘ilima. ‘Ilima is found from sea level to elevations > 1000 mas on multiple Hawaiian islands. Coastal populations are morphologically different from montane populations (Fig. 1). Coastal regions of Hawai’i are characterized by high sun exposure, warmer temperatures, high winds, salinity, and variation in water availability. Coastal populations of ‘ilima tend to be short and prostrate which likely helps them to withstand the windy environment (Fig. 1a). The leaves of these populations are covered on both surfaces in dense, soft hairs that give the leaves a silvery green appearance (Fig. 1b), which helps mitigate water loss by reflecting solar radiation, thereby lowering leaf temperature (Ehleringer and Björkman, 1978). Montane regions, on the other hand, provide very different challenges. Many other tall species grow on the slopes of these wet mountainous regions, which makes light competition a factor that plants may need to adapt to. Possibly due to this, montane populations are erect and shrub- or tree-like, capable of growing meters tall with strong, woody stems. These individuals have smooth, green foliage with serrated edges. Montane populations exhibit traits that may help them to compete for light availability. This montane morphology is not found in *S. fallax* populations on other Pacific Islands (Pejhanmehr, 2022).

**Figure 1:**
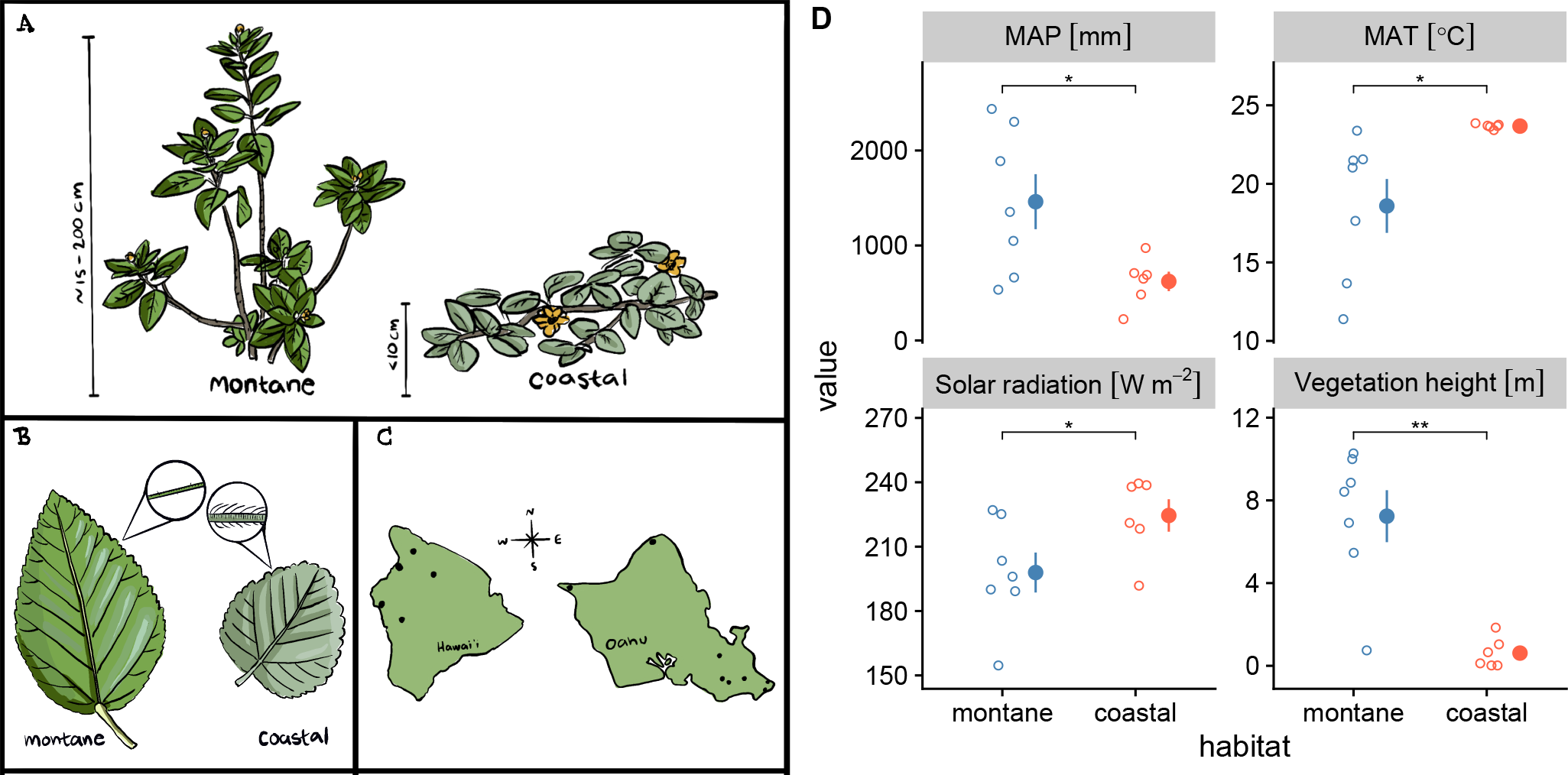
A. Typical growth form of montane (left) and coastal (right) ‘ilima plants and B. leaves. C. Map of the sites that were sampled on the islands of O’ahu and Hawai’i (aka Big Island). D. Climatic, light, and vegetation height comparisons between montane (blue) and coastal (orange) habitats sampled in this study. Open circles are values for the midpoint of each site transect; closed circles and intervals are the mean ± 1 standard error. The habitats differ significantly in mean annual precipitation (top-left), solar radiation (bottom-left), temperature (top-right), and vegetation height (bottom-right). MAP = mean annual precipitation; MAT = mean annual temperature; ns = not significant; * indicates 0.01 ≤ *P* < 0.05; ** indicates 0.001 ≤ *P* < 0.01.

Because of their contrasting habitat and morphology, we treat leaves from coastal and montane plants as representatives of sun and shade leaves, respectively, for testing hypotheses about amphistomy advantage. Specifically, the objectives of our study are to test whether 1) sun leaves of coastal ‘ilima plants will have greater AA than shade leaves of montane plants; and if so, is this because 2a) coastal plants have thicker leaves than montane plants and/or 2b) coastal plants have a *g*_smax,ratio_ closer to 0.5?

## MATERIALS AND METHODS

### Plant sampling and climate

We identified 7 suitable natural populations of ‘ilima on O’ahu and 5 on Hawai’i Island by consulting Yorkston and Daehler (2006) and citizen scientist records on iNaturalist (Anon, 2022) (Fig. 1c; Table 1). We avoided sites that appeared to be cultivated. We visited sites between August and November 2022. For logistical reasons, the sites on Hawai’i were sampled during one three-day trip. We haphazardly sampled eight plants distributed evenly between the highest and lowest elevation plants along a transect at each site. For safety and conservation reasons, transects were along a trail or road. We did not sample small individuals if there was risk removing leaves would cause mortality. From each plant, we collected two fully expanded leaves for trait measurements. We sampled stomatal traits on all leaves; leaf thickness on one leaf from three randomly selected plants per site; and, due to limited time, a single leaf from a single plant at the middle of each transect for gas exchange measurements. We downloaded climatic data on mean annual temperature, solar radiation, and vegetation height from the Climate and Solar Radiation of Hawai’i databases (Giambelluca et al., 2014) using the latitude and longitude at the middle of each transect. We also downloaded mean annual precipitation from 1978-2007 from the Rainfall Atlas of Hawai’i (Giambelluca et al., 2013). The spatial resolution is approximately 234 × 250m. The temperature data are calibrated from networks of meteorological stations operating in the late 20th century and 21st century; the solar radiation data are calibrated from satellite measurements collected between 2002 and 2009 (Giambelluca et al., 2014). We tested whether climatic variables differed among our coastal and montane populations using Welch’s two-sample *t*-test.

**Table 1:** ‘Ilima study site location information.

| Site | Island | Habitat | Latitude | Longitude | Elevation<br>(mas) |
| --- | --- | --- | --- | --- | --- |
| Kahuku Point | O‘ahu | coastal | 21.710 | -157.982 | 4 |
| Kaloko beach | O‘ahu | coastal | 21.293 | -157.661 | 4 |
| Kaloko-Honokōhau<br>national historical park | Hawai‘i | coastal | 19.676 | -156.024 | 6 |
| Ka‘ena Point | O‘ahu | coastal | 21.574 | -158.278 | 4 |
| Makapu‘u beach | O‘ahu | coastal | 21.313 | -157.661 | 3 |
| Puakō petroglyph park | Hawai‘i | coastal | 19.957 | -155.858 | 8 |
| Hawai‘i loa ridge | O‘ahu | montane | 21.294 | -157.727 | 352 |
| Hāloa ‘Āina | Hawai‘i | montane | 19.552 | -155.793 | 1567 |
| Ka‘ohe game<br>management area | Hawai‘i | montane | 19.817 | -155.616 | 1946 |
| Koai‘a tree sanctuary | Hawai‘i | montane | 20.048 | -155.737 | 970 |
| Mau‘umae Ridge | O‘ahu | montane | 21.305 | -157.779 | 248 |
| Wa‘ahila ridge | O‘ahu | montane | 21.314 | -157.793 | 357 |

### Leaf traits

#### Stomata

We estimated the stomatal density and size on ab- and adaxial leaf surfaces from all leaves. For pubescent leaves (usually coastal), we dried and pressed leaves for ≈ 1 week (Hill et al., 2014), carefully scraped trichomes off with a razor blade, and rehydrated the leaf. Rehydration restores leaf area to its fresh value (Blonder et al., 2012). For glabrous leaves, we used fresh leaves. We applied clear nail polish to both leaf surfaces of fresh or rehydrated leaves in the middle of the lamina away from major veins. After nail polish dried, we mounted impressions on a microscope slide using transparent tape (Mott and Michaelson, 1991). We digitized a portion of each leaf surface impression using a brightfield microscope (Leica DM2000, Wetzlar, Germany). We counted all stomata and divided by the visible leaf area (0.890 mm^2^) to estimate density and measured guard cell length from five randomly chosen stomata per field using ImageJ (Schneider et al., 2012).

#### Leaf thickness

We cut thin sections using two razor blades taped together. We sectioned the leaf in a petri dish of water, wet-mounted sections onto a slide, and took digital micrographs using a brightfield microscope, as described above. Leaf thickness is measured as the length from upper cuticle to lower cuticle.

#### Gas exchange measurements

At each site, we selected one representative leaf from one plant near the middle of the transect for gas exchange measurements using a portable infrared gas analyzer (LI-6800PF, LI-COR Biosciences, Lincoln, NE, USA). We estimated the photosynthetic rate (*A*) and stomatal conductance to water vapor (*g*_sw_) at saturating light (photosynthetic photon flux density (PPFD) = 2000 *μ*mol m^−2^ s^−1^), ambient CO_2_ (415 ppm), and *T*_leaf_ = 25.0–29.3°C. The midday irradiance in coastal ‘ilima typically meets or even exceeds a PPFD of 2000 *μ*mol m^−2^ s^−1^ and previous experiments with sun leaves revealed that 2000 *μ*mol m^−2^ s^−1^ is always at or near saturating irradiance. Even though lower irradiance may be saturating for montane leaves, we used this higher value for all leaves to standardize conditions.

We also estimated ‘amphistomy advantage’ (AA) *sensu* Parkhurst (1978), but with modification. For each leaf, we measured the photosynthetic rate of an untreated amphistomatous leaf (*A*_amphi_) over a range of *g*_sw_ values. We refer to this as an *A*–*g*_sw_ curve, which is described in more detail below. We compared the *A*–*g*_sw_ curve of the untreated leaf to the photosynthetic rate of pseudohypostomatous leaf (*A*_hypo_), which is the same leaf but with gas exchange through the upper surface blocked by a neutral density plastic (propafilm). Hypostomy refers to leaves with stomata only present on the lower, typically abaxial, surface. We refer to the untreated and partially blocked leaves as “amphi” and “pseudohypo”, respectively. AA is calculated as the log-response ratio of *A* compared at the same total *g*_sw_:

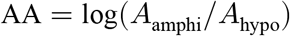

The log-response ratio is commonly used social and biological sciences (e.g. Hedges et al. (1999)). It is straightforward to interpret because values above 0 indicate a photosynthetic advantage of amphistomy, whereas values less than 0 indicate a disadvantage. The log-response ratio is preferable to the absolute difference because it indicates a proportional change in *A*, which facilitates comparisons across leaves and environments with different baseline photosynthetic rates. The irradiance of the light source in the pseudohypo leaf was higher because the propafilm reduces transmission. To compensate for reduced transmission, we increased incident PPFD for pseudohypo leaves by a factor 1/0.91, the inverse of the measured transmissivity of the propafilm. We also set the stomatal conductance ratio, for purposes of calculating boundary layer conductance, to 0 for pseudohypo leaves following manufacturer directions.

Fig. S1 illustrates our method for collecting *A*–*g*_sw_ curves. We collected two curves per leaf, an amphi (untreated) curve and a pseudohypo (treated) curve. To control for order effects, we alternated between starting with amphi or pseudohypo leaf measurements, though we did not detect an effect of treatment order on AA (results not shown). In the field, we acclimated the focal leaf to saturating light and high relative humidity (RH = 70%), as described above, until *A* and *g*_sw_ reach their maximum. We used these data as our estimates of maximum *A* and *g*_sw_. After that, we decreased RH to ≈ 10% to induce rapid stomatal closure without biochemical downregulation. Hence, *A*_amphi_ and *A*_hypo_ were both measured at low chamber humidity after the leaf had acclimated to high humidity. All other environmental conditions in the leaf chamber remained the same. We logged data until *g*_sw_ reached its nadir. We then repeated the process of acclimating the leaf to 70% RH and inducing stomatal closure with low RH with the other treatment (amphi or pseudohypo).

### Data analysis

#### Objective 1: Do coastal leaves have greater amphistomy advantage than montane leaves?

It is not feasible to record *A*_amphi_ and *A*_hypo_ at the exact same *g*_sw_. To overcome this, we fit *A*–*g*_sw_ curves using a linear regression of log(*g*_sw_) on *A* to interpolate modeled *A* for amphi and pseudohypo leaves at the same *g*_sw_. Let 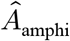 and 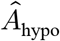 be the estimated *A* of the amphi and pseudohypo leaves, respectively. We estimated these quantities at the same *g*_sw_ using fitted parameters 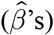:

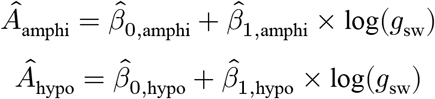

In 10 of 12 leaves, the minimum *g*_sw_ of the amphi curve was smaller than the maximum *g*_sw_ of the pseudohypo curve (i.e. the curves overlapped for a range of *g*_sw_ values). In those cases, we estimated 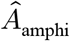 and 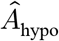 at the *g*_sw_ value in the middle of the range of overlap between the curves. In 2 of 12 leaves, the *A*–*g*_sw_ curves did not quite overlap because the minimum *g*_sw_ of the amphi curve was slightly greater than the maximum *g*_sw_ of the pseudohypo curve. In those cases, we estimated AA by extrapolating slightly, 1.98 × 10^−3^ and 3.29 × 10^−3^ mol m^−2^ s^−1^, beyond the measured curves to the *g*_sw_ value in between the curves. The vertical lines in Fig. S2 show the *g*_sw_ for each leaf. We estimated AA from 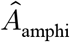 and 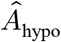 for each leaf using the log-response ratio shown above.

To estimate 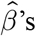 from the *A*–*g*_sw_ curve for each leaf, we fit Bayesian regressions using the *R* package **brms** version 2.20.4 (Bürkner, 2017) with MCMC sampling in *Stan* (Stan Development Team, 2023). We used CmdStan version 2.33.1 and **cmdstanr** version 0.6.1 (Gabry and Češnovar, 2023) to interface with *R* version 4.3.1 (R Core Team, 2023). We sampled the posterior distribution from 4 chains with 1000 iterations each after 1000 warmup iterations per chain. We estimated parameters and confidence intervals as the median and 95% quantile intervals of the posterior, respectively. The key prediction is that AA_coastal_ > AA_montane_, meaning the 95% confidence intervals of AA_coastal_ − AA_montane_ should be positive and not encompass 0.

#### Objective 2a: Are coastal leaves thicker than montane leaves?

We tested whether leaf thickness (log-transformed) varied between coastal and montane populations and among individuals within populations using a Bayesian mixed-effects model with habitat as a fixed effect and individual plant and site as random effects. We used the *R* package **brms** version 2.20.4 (Bürkner, 2017) to fit the model in *Stan* (Stan Development Team, 2023) with CmdStan version 2.33.1 and **cmdstanr** version 0.6.1 (Gabry and Češnovar, 2023). We sampled the posterior distribution from 4 chains with 1000 iterations each after 1000 warmup iterations per chain. We estimated the relationship between population average leaf thickness and AA measured from a single individual per population. We used this approach because most of the variation in leaf thickness was among sites and the plant selected for gas exchange measurements was not always among the plants randomly selected for leaf thickness, precluding individual level correlation. We propagated uncertainty about in AA and leaf thickness estimates by integrating over the entire posterior distribution sample for each variable. The key prediction is that the effect of leaf thickness on AA is positive, meaning the 95% confidence interval of the slope should be positive and not encompass 0.

#### Objective 2b: Is g_smax,ratio_ closer to 0.5 in coastal leaves than montane leaves?

We tested whether *g*_smax,ratio_ varied between coastal and montane populations and among individuals within populations using a Bayesian mutliresponse, mixed-effects model. The modeled response variables are stomatal count and guard cell length on each surface. Counts were modeled as negative binomially distributed variable from a latent stomatal density and a parameter *ϕ* to estimate overdispersion in counts relative to a Poisson model. For all traits, the explanatory variables were habitat as a fixed effect and leaf within individual plant, individual plant, and site as random effects. We used the *R* package **brms** version 2.20.4 (Bürkner, 2017) to fit the model in *Stan* (Stan Development Team, 2023) with CmdStan version 2.33.1 and **cmdstanr** version 0.6.1 (Gabry and Češnovar, 2023). We interpolated missing adaxial guard cell lengths from 6 out of 185 samples with zero adaxial stomata using the “mi” function in **brms** package. We sampled the posterior distribution from 4 chains with 1000 iterations each after 1000 warmup iterations per chain. From each posterior sample, we calculated *g*_smax,ratio_ as:

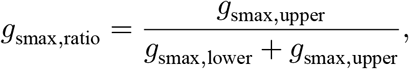

where *g*_smax,lower_ and *g*_smax,upper_ are maximum stomatal conductance to water vapor at *T*_leaf_ = 25° C on the lower and upper surface, respectively. The maximum stomatal conductance was calculated from stomatal density and length, assuming that stomata are fully open, following Sack and Buckley (2016):

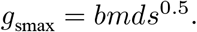

In this equation, *b* is a biophysical constant, *m* is a morphological constant, *d* is the stomatal density, and *s* is the stomatal complex area. We assume that *b*, which is determined by the molecular species, temperature, and air pressure, is the same for both surfaces; we assume that *m*, which is determined by guard cell allometry is also the same for both surfaces. Hence, the *b* and *m* constants cancel out of *g*_smax,ratio_ and only density and length (*l*), which is proportional to the square root of area, affect the ratio: *g*_smax_ ∝ *dl*.

We estimated the relationship between leaf *g*_smax,ratio_ and AA measured from a single leaf per population. We propagated uncertainty about AA and *g*_smax,ratio_ by integrating over the entire posterior distribution sample for each variable. The key prediction is that the effect of *g*_smax,ratio_ on AA is positive until *g*_smax,ratio_ < 0.5, meaning the 95% confidence interval of the slope should be positive and not encompass 0 in the domain *g*_smax,ratio_ < 0.5.

## RESULTS

Coastal ‘ilima are surrounded by shorter vegetation than their montane counterparts (Fig. 1d; Welch Two Sample *t*-test, *t*_6.67_ = 5.13, *P* = 0.002). The montane site with the lowest vegetation height is a remnant dry forest (Koai’a tree sanctuary) in a matrix of cattle pasture, hence the satellite derived vegetation height may be lower than what existed prior to human disturbance. Coastal sites receive greater average solar radiation at the top of the canopy (Fig. 1d; Welch Two Sample *t*-test, *t*_10.86_ = −2.22, *P* = 0.049); coastal sites are significantly warmer (Fig. 1d; Welch Two Sample *t*-test, *t*_6.01_ = −2.96, *P* = 0.025); and coastal sites receive less precipitation (Fig. 1d; Welch Two Sample *t*-test, *t*_7.45_ = 2.73, *P* = 0.028).

### Amphistomy advantage is greater in coastal leaves

Amphistomy increases photosynthesis in leaves of coastal ‘ilima plants more than that of montane plants. AA was significantly greater than 0 (95% confidence intervals did not overlap 0) in 5 of 6 coastal leaves, but only 1 of 6 montane leaves (Fig. 2; see Fig. S2 for individual curves). Overall, the average AA among coastal and montane leaves is 0.12 [0.077–0.15] and 0.027 [−0.0034–0.057], respectively; the difference in average AA between habitat types is AA_coastal_ −AA_montane_ = 0.09 [0.039–0.14]. Posterior predictions closely match observed values of *A* (Fig. S3), indicating an adequate model fit from which we can interpolate between measurements reliably. It also suggests that slight extrapolation beyond the data should be reliable, but this is less certain. When we remove two leaves where we extrapolated slightly beyond fitted *A*–*g*_sw_ curves, we estimate that AA_coastal_ is still positive, 0.081 [0.023–0.13], but the difference between coastal and montane leaves is smaller, 0.053 [−0.012–0.12], and confidence intervals slightly overlap 0. Maximum photosynthetic rate was slightly, but not significantly higher in coastal leaves (Welch Two Sample *t*-test, *t*_9.65_ = 1.6, *P* = 0.14); total stomatal conductance was similar (Welch Two Sample *t*-test, *t*_9.71_ = −0.09, *P* = 0.93) in coastal and montane leaves (Fig. S4). Water-use efficiency (*A*/*g*_sw_) was significantly higher in coastal leaves (Welch Two Sample *t*-test, *t*_9.99_ = 2.54, *P* = 0.03).

**Figure 2:**
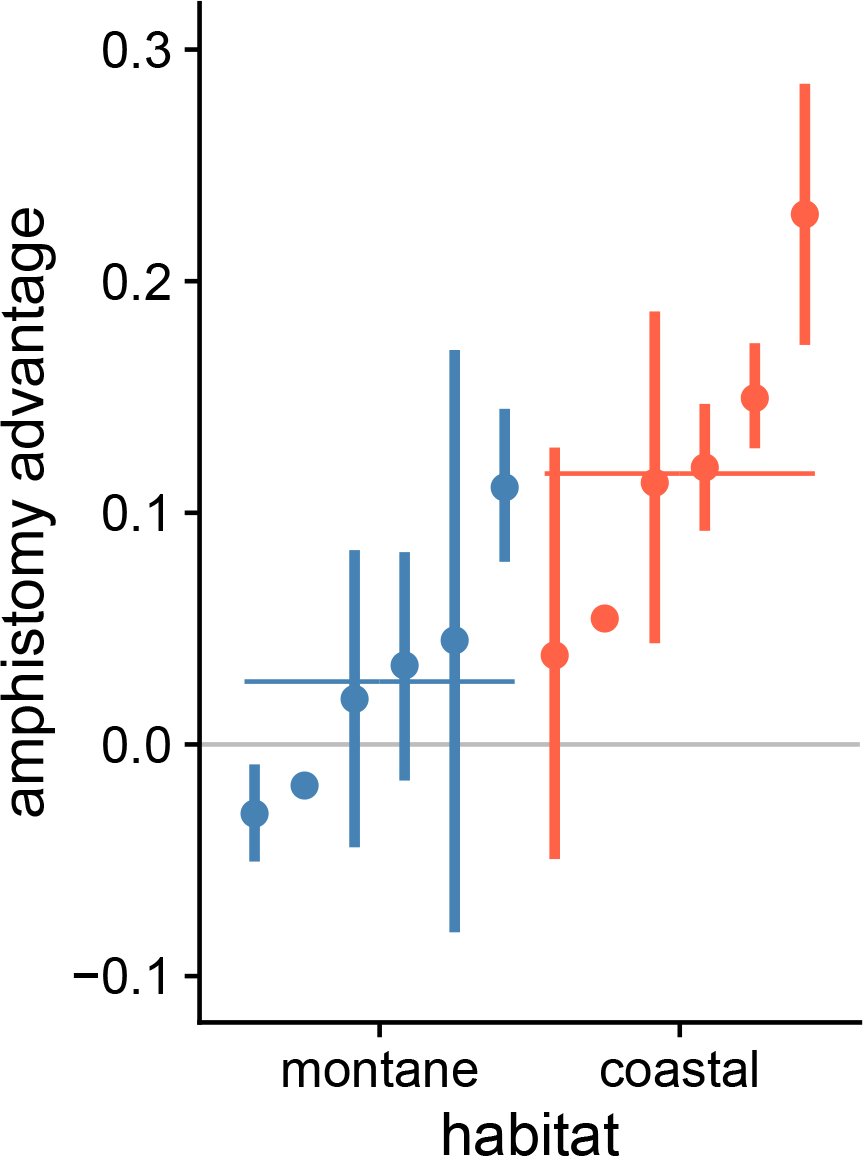
Coastal leaves benefit more from amphistomy than montane leaves. A positive amphistomy advantage (*y*-axis) means that the photosynthetic rate of an amphistomatous leaf is greater than that of an identical pseudohypostomatous leaf at the same overall *g*_sw_. Each pointinterval is the median posterior estimate plus 95% confidence interval of amphistomy advantage for that leaf. Each leaf is from a different montane (blue) or coastal (orange) site, arranged by habitat and ascending amphistomy advantage within habitat. The longer horizonal bars are the average amphistomy advantage for montane and coastal leaves. *g*_sw_, stomatal conductance to water vapor.

### Leaf thickness is associated with amphistomy advantage between but not within habitats

Coastal ‘ilima leaves are 91 [26–164] *μ*m thicker than their montane counterparts. Although coastal leaves are thicker and have greater AA, there is little relationship between leaf thickness and AA within habitats (Fig. 3A; slope = −0.11 [−0.28–0.035]).

**Figure 3:**
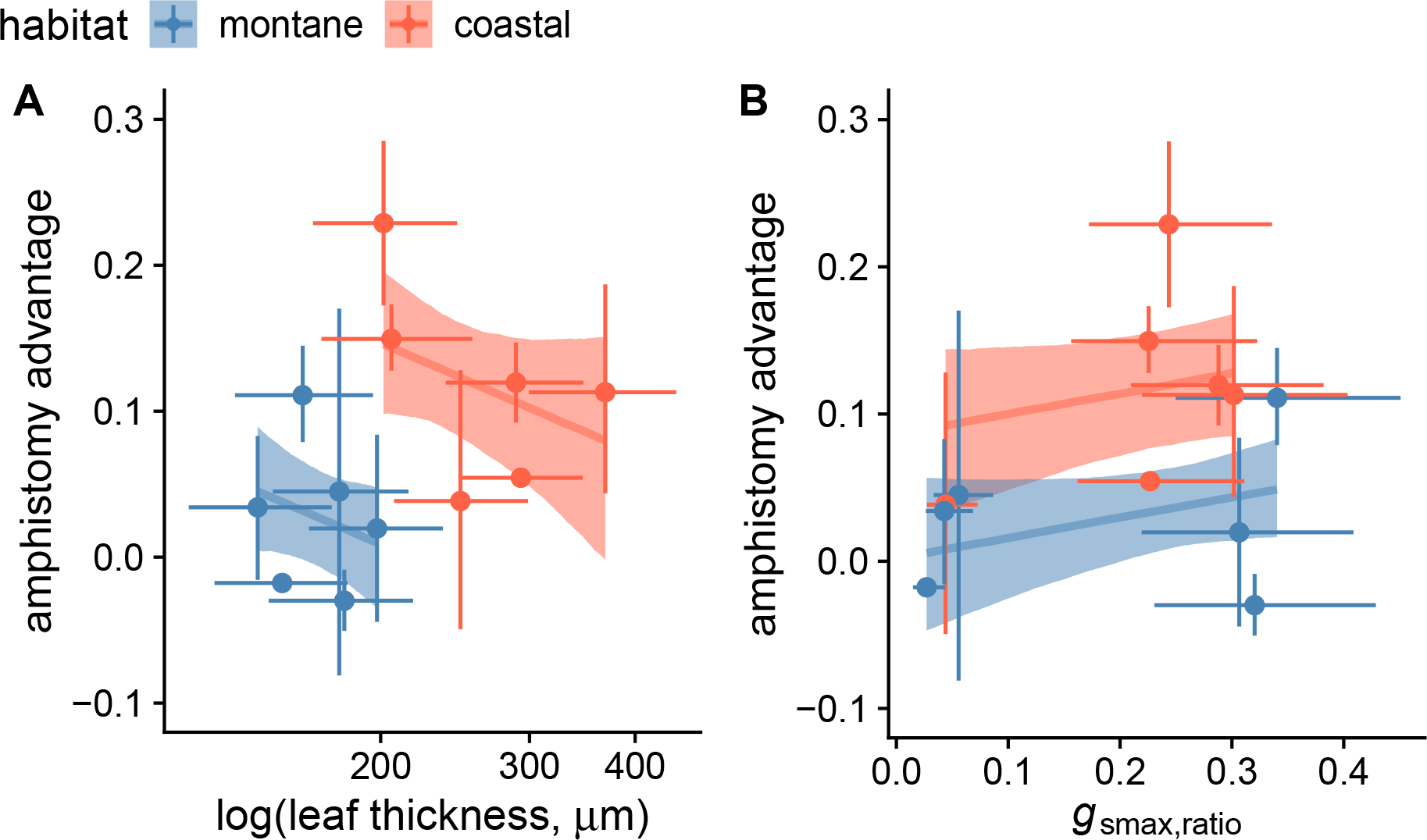
Relationships between leaf amphistomy advantage, (A) leaf thickness and (B) *g*_smax,ratio_ among ‘ilima (*Sida fallax*) plants from montane (blue) and coastal (orange) habitats in Hawai’i. A positive amphistomy advantage (*y*-axis) means that the photosynthetic rate of an amphistomatous leaf is greater than that of an identical pseudohypostomatous leaf at the same overall *g*_sw_. Each point-interval is the median posterior estimate plus 95% confidence interval of the trait value. Each leaf is from a different montane (blue) or coastal (orange) site. Lines are the estimated linear regression of (A) log(leaf thickness) and (B) *g*_smax,ratio_ on amphistomy advantage; ribbons are the 95% confident bands of the regression. Symbols: *g*_smax,ratio_, anatomical maximum stomatal conductance ratio; *g*_sw_, stomatal conductance to water vapor.

### *g*_smax,ratio_ is not associated with amphistomy advantage

Coastal and montane leaves have similar average *g*_smax,ratio_, the ratio of adaxial (upper) to total *g*_smax_, the anatomical maximum stomatal conductance to water vapor (Fig. S5); coastal leaves have 0.059 [−0.14–0.28] higher *g*_smax,ratio_ than montane leaves, but the 95% confidence intervals overlap 0 difference. The *g*_smax,ratio_ is somewhat bimodal among sites. Some sites in both habitats have leaves with *g*_smax,ratio_ < 0.07 and others with *g*_smax,ratio_ > 0.2 (Fig. S5). This is particularly noticeable in montane sites where those on the Big Island of Hawai’i all have low *g*_smax,ratio_ whereas those on O’ahu have relatively high *g*_smax,ratio_. There is no relationship between *g*_smax,ratio_ and AA in either habitat (Fig. 3B; slope = 0.14 [−0.057–0.34]) in our sample.

## DISCUSSION

Amphistomy is a seemingly simple way that leaves can increase carbon gain without significant additional water loss, yet it is rare in nature and we do not know why. The strong association between amphiostomy and sunny, open habitats suggests that amphistomy may benefit sun leaves more than shade leaves, but progress has been limited by the lack of evidence that amphiostomy *per se* improves photosynthesis in a given leaf. By experimentally blocking gas exchange through the upper surface in a controlled environment, we directly compared an amphistomatous leaf to an otherwise identical pseudohypostomatous leaf. This allows us to quantity the amphistomy advantage (AA) holding all else constant. Taking advantage of the steep climatic gradients in the Hawaiian archipelago, we applied this new method to show for the first time that sun leaves benefit 4.04 times more from amphistomy than shade leaves on ‘ilima (*Sida fallax*) plants (AA_coastal_ = 0.12 vs. AA_montane_ = 0.027). Coastal and montane ‘ilima leaves are likely good representatives of classic sun and shade leaf syndromes because 1) they vary in traits like reflective pubescence (Ehleringer and Björkman, 1978) and leaf thickness (Terashima et al., 2001) that typically characterize sun-shade adaptation; and 2) since ‘ilima shrubs are typically < 1m tall, they are shaded by trees in montane, but not coastal habitats (Fig. 1d). While this result has not yet been validated in other species, our results indicate that part of the reason amphistomatous leaves are found most commonly in high light habitats is that the adaptive benefit is greater in such environments.

If AA is typically greater in sun leaves than shade leaves, it could partially explain the distribution of amphi- and hypostoamtous leaves, but the precise mechanism(s) require further study. One hypothesis is that the internal airspace conductance, *g*_ias_, from stomata to mesophyll cell walls is lower in thicker sun leaves (Parkhurst, 1978). All else being equal, a leaf with lower *g*_ias_ will benefit more from amphistomy. Our results partially support this hypothesis. Coastal ‘ilima leaves with high AA (Fig. 2) are thicker than montane leaves, but the relationship between AA and leaf thickness within habitats is actually slightly negative (Fig. 3a), opposite our prediction. Since coastal and montane leaves differ in many respects besides thickness, we do not have enough data to conclude that leaf thickness explains the variation in AA between habitats. Alternatively, other biochemical or anatomical differences between coastal and montane leaves may explain why AA is greater in coastal leaves. The negative relationship, albeit nonsignificant in that 95% confidence intervals encompassed 0, between leaf thickness and AA could be explained if thicker leaves compensated by having a more porous mesophyll and/or less tortuous airspaces (Théroux-Rancourt et al., 2021).

A second natural hypothesis is that amphistomatous leaves with few adaxial (upper) stomata benefit less than those with similar densities on both surfaces. We predicted that leaves with *g*_smax,ratio_ closer to 0.5 would have higher AA based on biophysical models (Gutschick, 1984a). The logic is that a small number of stomata on the upper surface are insufficient to supply the entire upper mesophyll due to limited lateral diffusion (Morison et al., 2005). Our results do not support this hypothesis. Montane leaves from Big Island sites had low *g*_smax,ratio_ and low AA whereas low montane leaves on O’ahu had high *g*_smax,ratio_, but similarly low AA (Fig. 3b). Among coastal sites, the site with the lowest *g*_smax,ratio_ had the lowest AA, but there was little variation in *g*_smax,ratio_ among coastal leaves in our sample. We therefore cannot rule out that a larger sample of coastal leaves with greater variance in *g*_smax,ratio_ might support this hypothesis.

Two major implications from our study are that 1) photosynthesis in hypostomatous leaves is likely limited by CO_2_ concentration drawdown within leaf airspaces; and 2) amphistomy *per se* contributes to, but is not wholly responsible for, higher photosynthetic rates among amphistomatous leaves. The amphistomy advantage we observe in ‘ilima leaves implies decreased CO_2_ supply in pseudohypostomatous leaves because of concentration drawdowns in the leaf airspace. Limited diffusion through the airspace has long been hypothesized to depress photosynthesis in hypostomatous leaves (Parkhurst, 1994), with empirical support from helox studies (Parkhurst and Mott, 1990). However, these studies relied on interspecific comparisons of amphi- and hypostomatous leaves that differ systematically in many traits that affect gas exchange and photosynthesis (Xiong and Flexas, 2020). Our experimental approach overcomes this limitation and implies that the drop in CO_2_ concentration from substomatal cavities to the upper surface depresses photosynthesis.

Among land plants grown in a common garden, amphistomatous leaves have on average nearly 2× higher area-based photosynthetic rates (Xiong and Flexas, 2020), naively implying an AA ≈ log 2 = 0.69. This is much higher than our estimate of 0.12 among coastal ‘ilima leaves. The most likely explanation is that amphistomy is not the only cause of high photosynthetic rate. Indeed, species adapted to open, high light habitats with amphistomatous leaves also have higher concentrations of Rubisco, overall stomatal conductance, and photosynthetic capacity (Smith et al., 1997; Xiong and Flexas, 2020). For a leaf with high photosynthetic capacity that is well illuminated and hydrated, the major limitation becomes CO_2_. Under these conditions, amphistomy may substantially increase photosynthesis, as we observe in coastal ‘ilima leaves. Selection on increased photosynthesis under similar conditions may explain why crop leaves tend to increase stomatal density ratio during domestication (Milla et al., 2013).

Three limitations of this study are the small sample size, experimental design that precludes distinguishing genetic from environmental differences in leaf traits, and potentially confounding effects of other environmental differences besides light environment. Understanding the mechanistic basis of higher AA in sun leaves would require much larger sample sizes. Sun leaves tend to be thicker, more densely packed with mesophyll cells, and have greater photosynthetic capacity and higher stomatal conductance, among other traits (Lambers et al., 2008). Each of these factors and others potentially modulate AA. Quantifying the contribution of all these factors requires larger samples and additional measurements that are beyond the scope of this study, but exciting avenues for future research on leaf structure-function relations. Although many morphological traits that distinguish coastal and montane ‘ilima populations persist in a common environment (Yorkston and Daehler, 2006), we cannot distinguish between genetic effects and plastic responses to habitat as causes of difference in AA because we measured naturally occurring plants *in situ*. While disentangling genetic and plastic contributions is not necessarily important for understanding the distribution of amphistomatous leaves, it would be insightful to know about genetic and environmental contributions to trait variation. A reciprocal transplant would be able to determine the genetic and environmental contributions, as well their interaction, to trait variance in nature. However, reciprocal transplants cannot control for other differences between coastal and montane habitats besides vegetation height, such as temperature and precipitaiton. Experimental studies in controlled environments will be necessary to isolate the effects of light quantity and quality on AA.

## CONCLUSIONS

This study reports the first direct experimental evidence that having stomata open on both leaf surfaces, amphistomy, increases photosynthesis for a given total stomatal conductance, particularly in leaves from the type of open, sunny habitats where this trait is most common. By developing a straightforward experimental method to block gas exchange through the upper surface, we directly compared the photosynthetic rate of a leaf with gas exchange through both surfaces or just one, holding all other factors constant. In doing so, we found that coastal leaves of the indigenous Hawaiian ‘ilima (*Sida fallax*) enjoyed a greater photosynthetic benefit from amphistomy than nearby montane leaves living in more closed forest. This is not because amphistomatous leaves necessarily have greater leaf surface available for stomata, although that likely influences realized photosynthetic rates in natural populations. Rather, our experiments show that coastal amphistomatous leaves with the same total leaf stomatal conductance photosynthesize more than identical hypostomatous leaves. We cannot yet ascribe the difference in amphistomy advantage between coastal and montane leaves to particular physiological or anatomical variation, but this is a promising area for future research.

## Acknowledgments

The authors thank Kasey Barton and two anonymous reviewers for feedback on an earlier version of this manuscript, the University of Hawai’i honors council for guidance to GT, Tawn Speetjens for access to one montane site, TM Perez for advice on leaf sectioning, startup funds from the University of Hawai’i, NSF Award 1929167 to CDM, and NSF Award 2307341 to TNB.

## Author Contributions

GT and CDM contributed equally to all stages of this project; TNB contributed to development of the method and helped edit the manuscript.

## Data Availability Statement

Custom scripts are available on a GitHub repository (https://github.com/cdmuir/stomata-ilima) and will be archived on Zenodo with a DOI and stable URL upon publication. Raw data will be deposited on Dryad with a DOI and stable URL upon publication.

## Supporting Information

Additional supporting information may be found online in the Supporting Information section at the end of the article.

- Appendix S1: Supplemental figures and table

