## Supporting Information for "Amphistomy increases leaf photosynthesis more in coastal than montane plants of Hawaiian ‘ilima (*Sida fallax*)"

Table S1: Average traits values among ʻilima leaves at each site. $SD_{abaxial}$ is the stomatal density per $mm^2$ on the abaxial (lower) surface; $SD_{adaxial}$ is the stomatal density per $mm^2$ on the adaxial (upper) surface; $GCL_{abaxial}$ is the guard cell length in $\mu$m on the abaxial (lower) surface; $GCL_{adaxial}$ is the guard cell length in $\mu$m on the adaxial (upper) surface; Leaf thickness is the length from upper cuticle to lower cuticle in $\mu$m; $A$ is the photosynthetic rate in $\mu$mol $CO_2$ $m^{-2}$ $s^{-1}$; $g_{sw}$ is the stomatal conductance to water vapor in mol $m^{-2}$ $s^{-1}$.

| Site | Island | Habitat | $SD_{abaxial}$ | $SD_{adaxial}$ | $GCL_{abaxial}$ | $GCL_{adaxial}$ | Leaf thickness | $A$ | $g_{sw}$ |
|---|---|---|---|---|---|---|---|---|---|
| Kaloko-Honokōhau national historical park | Hawaiʻi | coastal | 412.99 | 92.31 | 14.73 | 20.32 | 189.60 | 17.40 | 0.123 |
| Puakō petroglyph park | Hawaiʻi | coastal | 310.12 | 52.55 | 13.35 | 20.94 | 194.20 | 17.85 | 0.134 |
| Kahuku Point | Oʻahu | coastal | 420.43 | 80.80 | 16.23 | 22.77 | 304.24 | 20.88 | 0.155 |
| Kaloko beach | Oʻahu | coastal | 400.27 | 109.03 | 15.69 | 23.26 | 399.54 | 31.28 | 0.314 |
| Kaʻena Point | Oʻahu | coastal | 370.14 | 120.34 | 17.61 | 22.89 | 295.82 | 34.77 | 0.336 |
| Makapuʻu beach | Oʻahu | coastal | 408.28 | 23.18 | 15.99 | 22.03 | 249.71 | 32.87 | 0.349 |
| Hāloa ʻĀina | Hawaiʻi | montane | 307.94 | 7.61 | 16.49 | 22.59 | 149.41 | 13.07 | 0.116 |
| Kaʻohe game management area | Hawaiʻi | montane | 270.24 | 13.07 | 14.87 | 21.51 | 183.72 | 12.94 | 0.130 |
| Koaiʻa tree sanctuary | Hawaiʻi | montane | 318.58 | 9.55 | 13.89 | 22.86 | 138.89 | 27.88 | 0.358 |
| Hawaiʻi loa ridge | Oʻahu | montane | 329.25 | 115.56 | 17.83 | 21.76 | 205.97 | 21.95 | 0.215 |
| Mauʻumae Ridge | Oʻahu | montane | 298.77 | 138.32 | 16.53 | 20.72 | 162.02 | 24.60 | 0.436 |
| Waʻahila ridge | Oʻahu | montane | 346.75 | 150.26 | 17.76 | 20.61 | 193.33 | 13.96 | 0.194 |

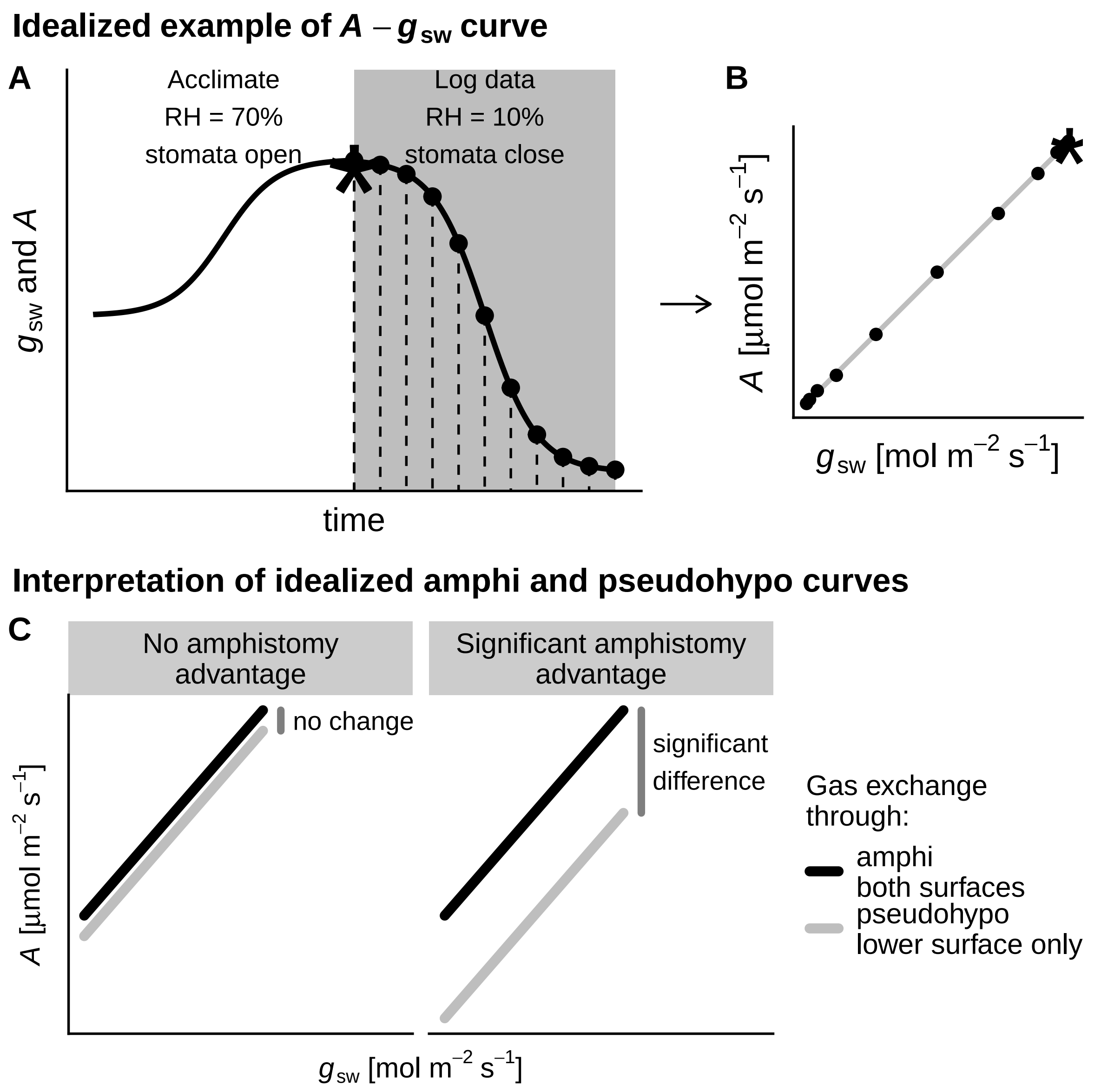


Figure S1: Idealized method for collecting $A$–$g_{sw}$ curves on either amphi or pseudohypo leaves. A) After clamping the leaf into the LI-6800 chamber, it acclimates to high light (PPFD $=$ 2000 $\mu$mol m$^{-2}$ s$^{-1}$) and humidity (RH $= 70\%$). This induces stomata to open, increasing $g_{sw}$ and $A$ until they reach a maximum. We abruptly lower the chamber humidity to $\approx 10\%$ to close stomata and log data (black points) until $g_{sw}$ and $A$ reach their nadir. B) We fit $A$–$g_{sw}$ curves to logged data points. The asterisk in both panels indicates the data point used for maximum $A$ and $g_{sw}$. C) AA is low (left panel) when the photosynthetic rate of an amphi leaf is similar to a pseudohypo leaf at the same total $g_{sw}$ ($x$-axes); large AA (right panel) is indicated when an amphi leaf has a higher photosynthetic rate than a pseudohypo leaf. Abbreviations: $A$ is the photosynthetic rate; AA is the amphistomy advantage; $g_{sw}$ is the stomatal conductance to water vapor; PPFD is photosynthetic photon flux density; RH is relative humidity.

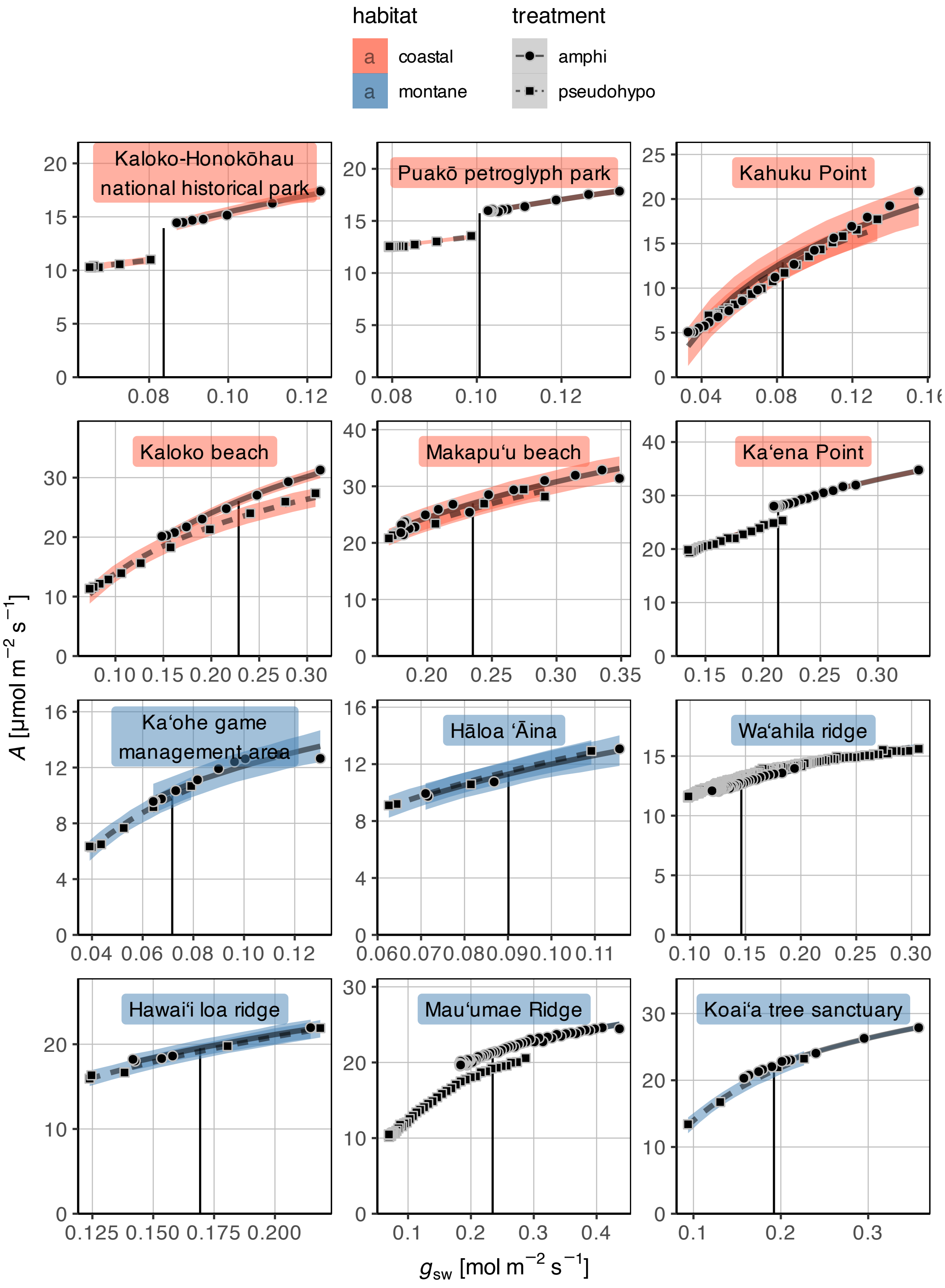


Figure S2: See next page.

Figure S2: (Continued from previous page.) Individual $A$–$g_{\text{sw}}$ curves used to estimate AA. For each leaf, one per site, we measured $A$ ($y$-axis) over a range of $g_{\text{sw}}$ ($x$-axis) on the same leaf with two treatments: 'amphi' (circles, solid line) leaves were untreated; 'pseudohypo' (squares, dashed line) leaves had no conductance through the upper (adaxial) surface. In all coastal (orange) and montane (blue) leaves, we fit generalized additive models and 95% confidence ribbons to estimate AA at a $g_{\text{sw}}$ where the curves overlap (vertical black line). In leaves from Kaloko-Honokōhau national historical park and Puakō petroglyph park, we extrapolated sligtly beyond fitted curves because they did not quite overlap. Symbols: AA, amphistomy advantage; $A$, photosynthetic rate in $\mu$mol $CO_2$ $m^{-2}$ $s^{-1}$; $g_{\text{sw}}$, stomatal conductance to water vapor in mol $m^{-2}$ $s^{-1}$.

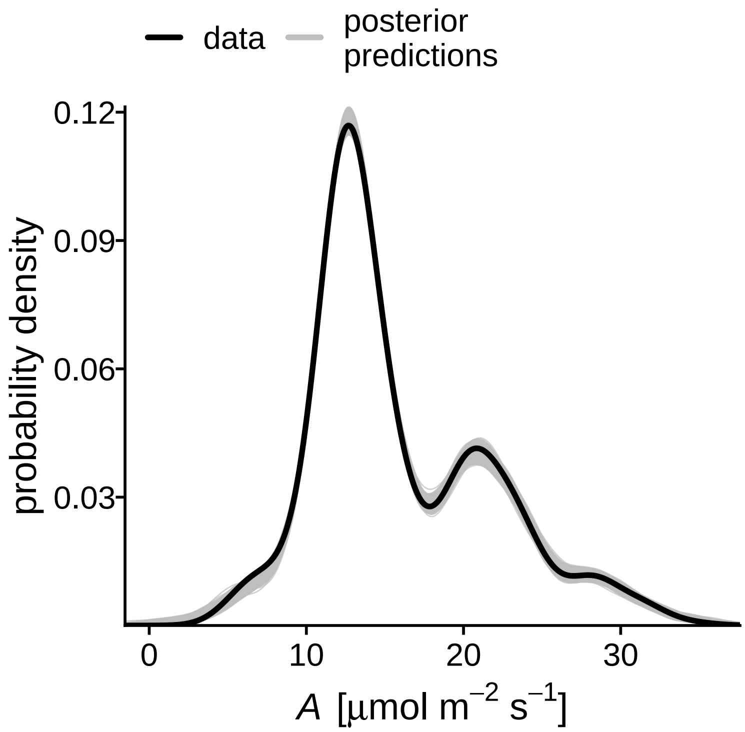


Figure S3: Posterior predictions (thin grey lines) from fitted $A$–$g_{\text{sw}}$ curves closely match the observed distribution (thick black line), indicating that the statistical model adequately captures variation in the response variable over the measured range. Symbols: $A$, photosynthetic rate in $\mu$mol $CO_2$ $m^{-2}$ $s^{-1}$; $g_{\text{sw}}$, stomatal conductance to water vapor in mol $m^{-2}$ $s^{-1}$.

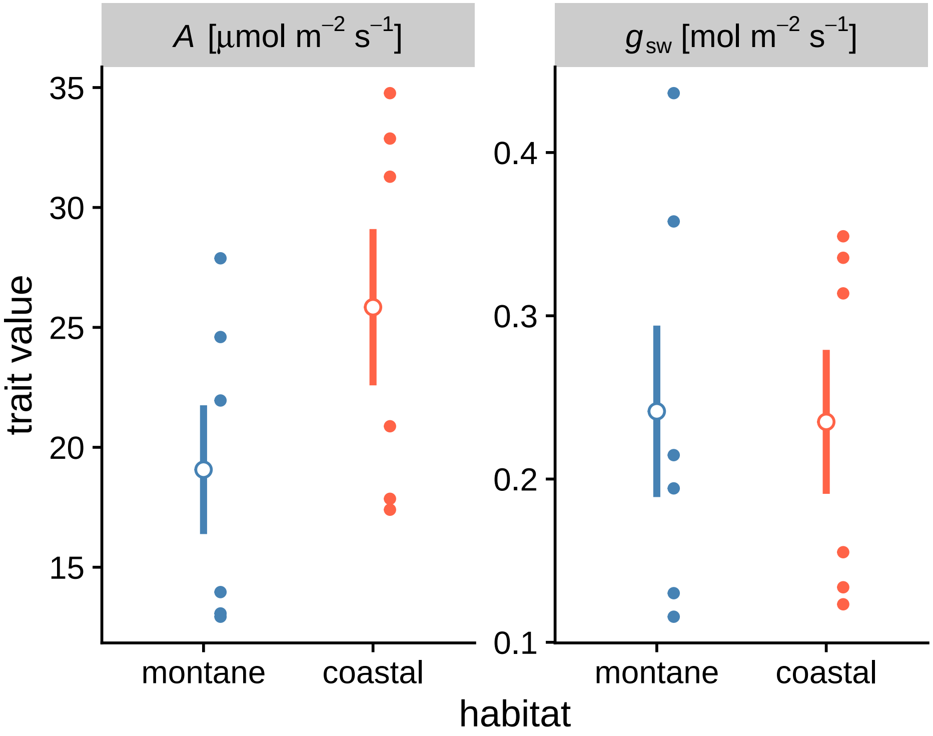


Figure S4: The photsynthetic rate (left facet) and stomatal conductance to water vapor (right facet) of montane (blue) and coastal (orange) ʻilima leaves. Each point-interval is the median posterior estimate plus 95% confidence interval of trait value for that habitat. Smaller points next to each point-interval are the $g_{\text{smax,ratio}}$ of individual plants, one per site. Symbols: $A$, photosynthetic rate in $\mu$mol $CO_2$ $m^{-2}$ $s^{-1}$; $g_{\text{sw}}$, stomatal conductance to water vapor in mol $m^{-2}$ $s^{-1}$.

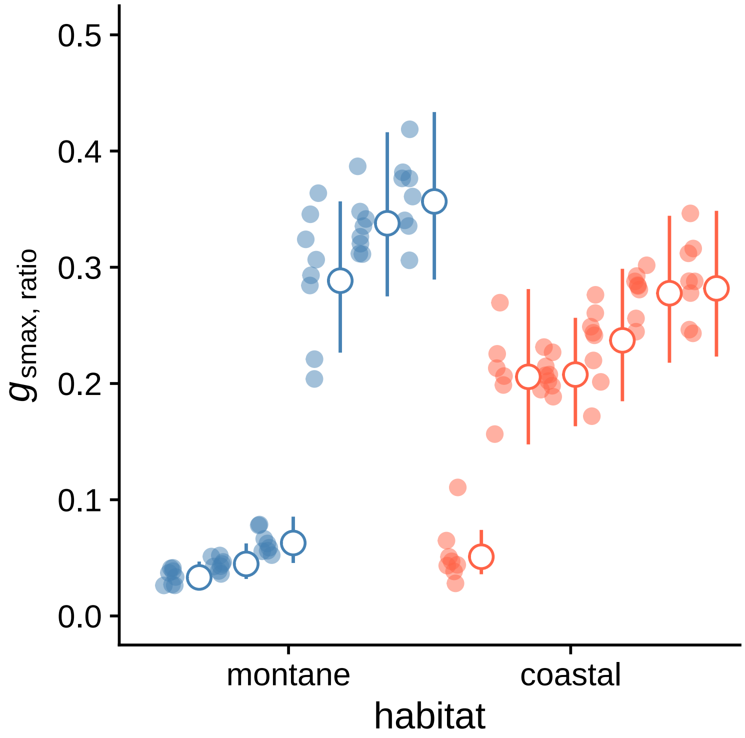


Figure S5: The $g_{\text{smax,ratio}}$ ($y$-axis) of montane (blue) and coastal (orange) ‘ilima leaves. Each point-interval is the median posterior estimate plus 95% confidence interval of $g_{\text{smax,ratio}}$ for that site. Sites are arranged by habitat and ascending $g_{\text{smax,ratio}}$ within habitat. Smaller, transparent points next to each point-interval are the $g_{\text{smax,ratio}}$ of individual plants. Symbols: $g_{\text{smax,ratio}}$, the ratio of anatomical maximum stomatal conductance to water vapor on the the adaxial surface over the total.
